# Application of the SMALP technology to the isolation of GPCRs from low-yielding cell lines

**DOI:** 10.1101/2021.01.26.428305

**Authors:** Daniele Tedesco, Maciej Maj, Paulina Malarczyk, Andrea Cingolani, Mirko Zaffagnini, Artur Wnorowski, Tiziana Benelli, Rita Mazzoni, Manuela Bartolini, Krzysztof Jóźwiak

**Affiliations:** Department of Pharmacy and Biotechnology, University of Bologna, via Belmeloro 6, Bologna, Italy; Institute for Organic Synthesis and Photoreactivity, National Research Center (ISOF-CNR), via Gobetti 101, Bologna, Italy; Department of Biopharmacy, Medical University of Lublin, ulica Chodźki 4a, Lublin, Poland; Department of Industrial Chemistry, University of Bologna, viale del Risorgimento 4, Bologna, Italy; Department of Pharmacy and Biotechnology, University of Bologna, via Irnerio 42, Bologna, Italy

**Keywords:** Membrane proteins, GPCR solubilization, styrene–maleic acid co-polymers, β_2_-adrenergic receptor, HEK293T cells

## Abstract

The ability of styrene–maleic acid (SMAc) co-polymers to spontaneously insert into biological membranes can be exploited to extract G protein-coupled receptors (GPCRs) embedded in styrene–maleic acid lipid particles (SMALPs), preserving the native environment around the protein and thus enhancing the feasibility of functional studies. So far, the SMALP technology has been primarily employed on non-mammalian cells and protocols are not optimized for adherent human cell lines, which cannot be harvested in large amounts. In this work, a fine investigation of key parameters affecting the formation of SMALPs was undertaken with the purpose of maximizing the yield of extraction of a recombinant form of human β_2_-adrenergic receptor (rhβ_2_AR) from HEK293T cells. The study highlighted an important influence of ionic strength on the membrane solubilization efficiency and GPCR purification yield of SMAc co-polymers: by lowering the salt concentration of all buffers used in previously published SMALP protocols, the water solubility and extraction efficiency of the selected SMAc co-polymer (commercially supplied as a potassium salt) were enhanced. In-line combination of size-exclusion chromatography (SEC) with immobilized metal affinity chromatography (IMAC) allowed further improvement of the final rhβ_2_AR yield by reducing the loss of SMALP-embedded GPCRs during the fractionation and purification of SMALPs. The overall findings of this study show that the available SMALP protocols can be significantly optimized in several aspects in order to increase the efficiency of GPCR solubilization and isolation from low-yielding expression systems.

**Highlights:**

- The SMALP technology enables a direct solubilization of GPCRs from cell membranes.
- The isolation of GPCRs from mammalian cells is usually a low-yield procedure.
- SMALPs embedding the β_2_-adrenergic receptor were prepared from a HEK293T cell line.
- The ionic strength of buffers plays a key role in SMALP formation and isolation.
- Standard SMALP protocols can be finely optimized to increase purification yields.

**Graphical abstract:** 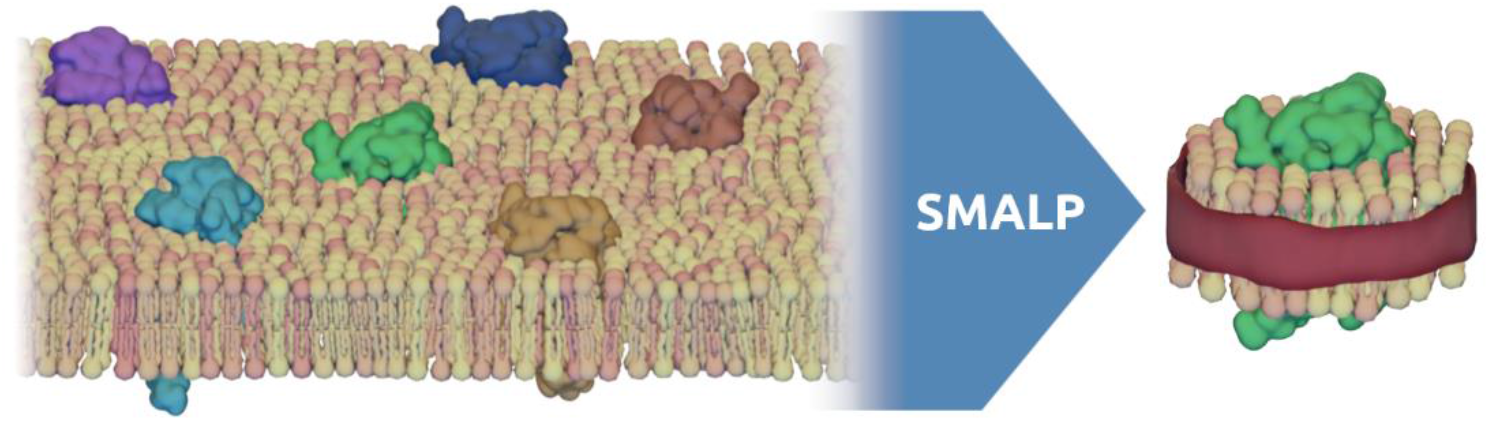

## 1 Introduction

The isolation of proteins from cell membranes is a process of paramount importance in the field of biochemistry. The feasibility of structural and functional studies on membrane proteins (MPs) relies on their genuine extraction from their natural environment and the solubilization of their native form in aqueous media. Several strategies have been developed for this specific purpose over the last five decades [1–3]. Human-derived MPs, especially G protein-coupled receptors (GPCRs), were shown to interact closely with the lipid constituents of biological membranes [4–7] and depend on these interactions for their function [8–10]: the most glaring example was the binding of cholesterol, an interaction known to greatly improve the crystallization of GPCRs [5,11]. MPs were also shown to be highly regulated by post-translational modifications (PTMs), which are organism- or, sometimes, even tissue-dependent. [12–14]. This complex network of molecular interactions is of key importance for the correct functioning of GPCRs. Therefore, the preservation of the native environment of biological membranes is a highly desirable condition for improving the significance of functional studies on MPs.

A new strategy for MP solubilization, based on the use of styrene–maleic acid (SMAc) co-polymers, has recently gathered increasing attention from the biochemistry community thanks to a combination of promising features [15]. The styrene–maleic acid lipid particle (SMALP) technology takes advantage of the amphiphilic properties of SMAc co-polymers, which can interact simultaneously with the hydrophobic core of lipid bilayers and with water. The result of such interactions is the formation of SMALPs, water-soluble disc-shaped lipid nanostructures embedding MPs and larger protein complexes, which can be obtained directly from crude membranes or even living cells with apparently minimal disruption of the native lipid environment around MPs. The ability of SMAc co-polymers to spontaneously insert into biological membranes can be exploited to avoid the use of surfactants for MP extraction, which may cause a loss in function through delipidation, and synthetic lipids membranes for mimicking the native environment of MPs [1,16]. The first published SMALP protocols are strikingly simple and straightforward, further increasing the appeal of this method for developing structural and functional studies on MPs in near-native conditions [15]. The apparent simplicity of this strategy depends on the properties of the cell membranes used for MP expression, on the structure of the employed SMAc co-polymer and on several environmental factors which may have a dramatic impact on the final yield of MP solubilization [17]. Most applications of the SMALP technology employ bacterial or insect expression systems, which can be grown in large quantities using suspension cultures, thus reducing the need for optimizing SMALP formation yields. Indeed, the use of human cell lines as a source of human GPCRs for functional studies poses important technical issues, such as low growth rates, high cost and limited scalability of adherent cell lines. Combined with the challenge of preserving the native membrane environment of human GPCRs, these issues can curtail the final yield of GPCR production, prompting the need for a careful examination of efficiency bottlenecks in the process of SMALP formation and purification.

The importance of environmental factors, such as pH, ionic strength and presence of divalent cations or temperature, in affecting the MP solubilization efficiency of SMAc is supported by increasing experimental evidence [17,18]. The effects of different polymerization strategies, styrene/maleic acid (Sty/MAc) ratios, average chain lengths and hydrolysis levels on the MP solubilization efficiency of SMAc co-polymers have also been described in the literature [19–21]. The ratio between the concentrations of membrane suspensions and SMAc co-polymer used in the preparation of SMALPs might also play a role, considering that an excess of SMAc can interfere with the downstream purification of MPs by affinity-based separation techniques. Finally, the composition of the biological membrane is a critical factor, as different SMAc co-polymers might display different MP solubilization efficiencies on different cell lines. All these factors become even more significant when dealing with small amounts of biological material, which is typically the case when working with human cell lines overexpressing GPCRs. There has been a limited number of published studies focused on the optimization of the MP solubilization efficiency of SMAc co-polymers for use with biological membranes, most of which dealt with artificial model membrane systems.

Herein, a recombinant form of the human β_2_-adrenergic receptor (rhβ_2_AR) was designed and transfected into a human cell line with the aim of generating SMALPs with an embedded GPCR surrounded by a native annular lipid shell. To overcome limitations related to the use of a human cell line, two commercially available SMAc co-polymers, namely Xiran SL 25010 P20 (X25, Sty/MAc ratio: 3:1, n/n) and Xiran SL 30010 P20 (X30, Sty/MAc ratio: 2.3:1, n/n), and an in-house synthesized SMAc co-polymer (C30, Sty/MAc ratio: 2.3:1, n/n) were compared in terms of MP solubilization efficiency. A fine optimization of the SMALP formation and isolation protocols was required to improve the yield of SMALP production; the outcomes of this investigation offer a panel of key experimental parameters to be taken into consideration when the SMALP technology needs to be applied to the solubilization of GPCRs starting from adherent human cell lines.

## 2 Materials and methods

### 2.1 Chemicals

Commercial styrene–maleic acid (SMAc) co-polymers X25 (Xiran SL 25010 P20, lot number PL0021601682-EH-A; 20.6% in water, m/m) and X30 (Xiran SL 30010 P20, lot number PL0021301582-EH-A, 22.3% in water, m/m) were kindly provided by Polyscope Polymers BV (Netherlands). For the synthesis of SMAc co-polymer C30, styrene (Sty), maleic anhydride (MAn), 2,2’-azobisisobutyronitrile (AIBN), tetrahydrofuran (THF), sodium hydroxide (NaOH), potassium hydroxide (KOH), anhydrous magnesium sulfate (MgSO_4_), chromium(III) acetylacetonate (Cr(acac)_3_), methanol (MeOH), chloroform (CHCl_3_), ethanol (EtOH, 99.9%) and deuterated solvents (D_2_O, acetone-d_6_ and DMSO-d_6_) were purchased from Sigma-Aldrich (Merck, USA). Commercial Sty was treated with a 5% (m/v) aqueous NaOH solution, washed with water, dried for 24 h on MgSO_4_ and distilled at 16 mmHg (*T*_b_ = 35 ° °C). AIBN was crystallized from EtOH before use. THF was purified and dried according to the reported procedures and stored under nitrogen [22]. MAn was used as received.

Buffers for the preparation of SMALPs were prepared in ultrapure water treated with a Milli-Q purification system (Merck, USA). Potassium phosphate monobasic (KH_2_PO_4_), potassium phosphate dibasic (K_2_HPO_4_), sodium chloride (NaCl), tris(hydroxymethyl)aminomethane (Tris), hydrochloric acid (HCl), glycerol, sucrose, calcium chloride (CaCl_2_) dihydrate, protease inhibitor cocktail tablets (SigmaFAST), ethylenediaminetetraacetic acid disodium salt (Na_2_EDTA), nickel(II) chloride (NiCl_2_) hexahydrate, imidazole and polyethylene glycol sorbitan monolaurate (Tween-20) were purchased from Sigma-Aldrich (Merck, USA).

The pX459 plasmid was purchased from Addgene (USA). Dulbecco’s Modified Eagle Medium (DMEM), fetal bovine serum (FBS), penicillin-streptomycin solution (5000 U/mL penicillin, 5000 μg/mL streptomycin), ampicillin sodium salt, the restriction endonuclease DpnI, the Turbofect cationic polymer-based transfection reagent, the GeneJet DNA spin column-based purification kit and the endotoxin binding reagent were purchased from ThermoFisher Scientific (USA). Puromycin solution (10 mg/mL) was purchased from Invivogen (France).

### 2.2 Production of recombinant human β_2_-adrenergic receptor (rhβ_2_AR)

#### 2.2.1 Plasmid construction

The employed rhβ_2_AR form was purposely designed for the small-scale production of SMALP-embedded GPCR samples. The gene encoding for the human β_2_-adrenergic receptor (rhβ_2_AR) was inserted into a pX459 plasmid flanked by appropriate tagging and signaling sequences: **(a)** a cleavable hemagglutinin signal sequence (ssHA) to enhance translocation and expression on the cell surface and a FLAG peptide tag on the *N*-terminal side; **(b)** the tobacco etch virus (TEV) endoprotease site, a 10×His peptide tag, a 12-residue spacer (SSGLPRGSHMAS) and a 6×His peptide tag on the *C*-terminal side (Figure 1). The gene insert was synthesized in whole by a commercial vendor (Integrated DNA Technologies, USA) and cloned into the pX459 backbone using Gibson assembly. Prior to cloning, the amplified backbone and insert DNA were cleaned up by digestion of template DNA using the restriction endonuclease DpnI and purified with a microcolumn-based system (Zymo Research, USA). The plasmid construct was transformed into an electrocompetent strain of *Escherichia coli* DH10β cells (ThermoFisher Scientific, USA) via electroporation; selection of clones was conducted with use of ampicillin (120 μg/mL), exploiting the ampicillin resistance gene (AmpR) present in the pX459 backbone. Bacterial clones were tested for the presence of the DNA insert by polymerase chain reaction (PCR) amplification and subsequent Sanger sequencing to verify the correct nucleotide sequence throughout the inserted fragment. Positive clones were selected and used for plasmid amplification; plasmids were then purified using the GeneJET purification kit with an addition of endotoxin binding reagent.

**Figure 1.**
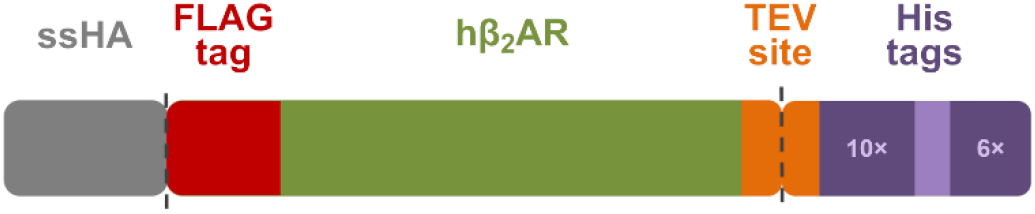
Schematic representation of the DNA construct for the expression of rhβ_2_AR; dashed lines denote cleavage sites.

#### 2.2.2 Cell transfection and culture

The human embryonic kidney 293T cell line (HEK293T; American Type Culture Collection, USA) was chosen as the mammalian system for rhβ_2_AR overexpression. Cells were cultured in DMEM supplemented with 10% FBS, penicillin (50 U/mL) and streptomycin (50 μg/mL). Cell lines were cultivated for a minimum of 3 and a maximum of 12 sub-culturing cycles before transfection. Transfection was performed using the Turbofect transfection reagent according to the manufacturer’s protocol. Transfection conditions were controlled by a means of parallel transfection with a control plasmid encoding far-red fluorescent protein mKate. Fluorescent microscope pictures of cells were obtained at 586 nm excitation and 646 nm emission wavelengths using a “FLoid” cell imaging station microscope system (ThermoFisher Scientific, USA) to confirm the correct transfection process and correct expression of the control protein. Subsequently, transfected cell cultures underwent selection using puromycin (1 μg/mL), exploiting the puromycin resistance gene (puroR) present in the pX459 backbone. Antibiotic selection was conducted for 21 days, thereon cells were moved to 15 cm diameter Petri dishes and grown up to ~80% confluency in 20 such dishes. Upon achieving desired confluence, cells were harvested mechanically using a cell scraper, suspended in cold PBS and centrifuged at 500×*g* for 5 min at 4 °C.

#### 2.4.3 Preparation of cell membrane suspensions

Harvested cells were lysed mechanically in lysis buffer LB (Tris-HCl 50 mM, sucrose 250 mM, CaCl_2_ 250 μM, pH 7.4 + SigmaFAST protease tablet) using a rotor-stator homogenizer (Pro Scientific, USA). The isolation of cell membranes from lysates was achieved by a two-step centrifugation protocol: a first centrifugation cycle was carried out at 750×*g* for 20 min at 4 °C, then the supernatant was ultracentrifuged at 100000×*g* for 20 min at 4 °C using an Optima L-100K ultracentrifuge (Beckman Coulter, USA) equipped with a SW 40 Ti swinging bucket rotor. The resulting cell membrane pellets were resuspended with suspension buffers (SB) of varying NaCl content (Table 1) in appropriate volumes to achieve the desired final concentration of membrane fractions.

**Table 1.** Composition of suspension buffers (SB) used for the preparation of cell membrane suspensions.

| Label | Buffer composition | pH |
| --- | --- | --- |
| SB0 | [Tris HCl 50 mM]/glycerol 90:10, v/v | 8.0 |
| SB250 | [Tris HCl 50 mM, NaCl 250 mM]/glycerol 90:10, v/v | 8.0 |
| SB500 | [Tris HCl 50 mM, NaCl 500 mM]/glycerol 90:10, v/v | 8.0 |

### 2.3 Synthesis and characterization of SMAc co-polymer C30

SMAc co-polymer C30 was obtained by hydrolysis of the corresponding styrene–maleic anhydride (SMAn) co-polymer N30 which was synthesized in a glass vial using AIBN as the radical initiator and dry THF as the solvent (total monomer concentration: 0.05 g/mL). The monomers (Sty/MAn ratio: 8:1, n/n), and AIBN (4%, n/n; in dry THF) were transferred into the vial under nitrogen atmosphere, submitted to several freeze–thaw cycles and heated at 60 °C for 17 h. The reaction was stopped by adding MeOH and the solvent was evaporated under reduced pressure. The resulting solid material (N30) was dissolved in CHCl_3_, precipitated in an excess of methanol, then dried under vacuum for several days up to constant weight. The co-polymer composition was determined by ^13^C-nuclear magnetic resonance (NMR). The hydrolysis of N30 to C30 was carried out in alkaline conditions, following a previously reported procedure [15] with slight modifications. In particular, the reaction time was increased up to 5 days and NaOH (1 M) was added several times. The reaction mixture was then cooled at room temperature and the insoluble part collected and discarded. The pH of the solution was then reduced to < 5 by addition of concentrated HCl to precipitate C30, which was washed three times with water and collected by centrifugation. The precipitate was then resuspended in NaOH (0.6 M) and the pH adjusted to pH 8 with HCl (2 M). Finally, the co-polymer was lyophilized using a freeze-dryer and characterized.

^1^H- and ^13^C-NMR spectra were carried out at 25 °C on an Inova 600 spectrometer (Varian, USA; ^1^H, 599.7 MHz; ^13^C, 150.8 MHz); chemical shifts were referenced internally to residual solvent peaks. The NMR spectra of N30 were obtained in acetone-d_6_ (100 mg of co-polymer in 0.6 mL of solvent) in the presence of Cr(acac)3 (0.1 M), while those of C30 were obtained in DMSO-d_6_ or D_2_O. Attenuated total reflectance Fourier-transform infrared (ATR-FTIR) spectra were recorded on an Alpha Platinum-ATR spectrophotometer (Bruker, Germany) equipped with an ATR Diamond window. The number- and weight-average molecular weights of the co-polymer (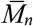 and 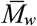) were determined in THF solution by size-exclusion chromatography (SEC), using a HPLC Lab Flow 2000 apparatus (Knauer, Germany) equipped with a 7725i injection valve (Rheodyne, USA), a Phenogel 5-μm MXL column (Phenomenex, Italy) and a RI K-2301 refractive index detector (Knauer, Germany). The calibration curve was obtained with standard monodisperse polystyrene samples.

#### SMAc co-polymer C30

*ATR-FTIR:* 3026 (*ν*_CH_, arom.), 2926 (*ν*_CH_, aliph.), 1556 (*ν*_COO^−^_), 1493, 1452 (*ν*_C=C_, arom.), 699 (*δ*_CH_, arom.) cm^−1^. ^*1*^*H-NMR (D*_*2*_*O):* 7.6−5.8 (CH, arom.), 2.9–0.4 (CH, CH_2_, aliph.) ppm. ^*13*^*C-NMR (D*_*2*_*O):* 181.8 (C(O)O), 153.7−135.7 (C_q_, arom.), 131.1−124.4 (CH, arom.), detectable peaks 49.8, 44.3, 40.3, 32.6 (CH, CH_2_, aliph.) ppm.

### 2.4 DLS assays on the solubility of SMAc co-polymers

Dynamic light scattering (DLS) measurements were carried out at 25 °C on a Zetasizer Nano ZS size analyzer (Malvern Instruments, UK) equipped with a He–Ne laser light source (633 nm wavelength), using disposable plastic UV micro-cuvettes with a 1 cm optical pathlength (Brand, Germany), non-invasive back scatter (NIBS) detection at a 173 ° angle, an equilibration time of 15 s and automatic positioning, attenuation and duration of measurements; data were averaged over 3 measurements with 0 s delay. The built-in values of refractive index and viscosity for water at 25 °C (*n* = 1.330; *η* = 0.8872 cP) were used as parameters for the dispersant. SMAc stock solutions (5%, m/v) were prepared in water for commercial co-polymers X25 and X30, and in KOH (0.15 M in water) for the synthesized co-polymer C30. DLS samples were then prepared in water, in potassium phosphate buffers (KP) of varying pH and salt concentrations (Table 2), and in suspension buffer SB500 ([Tris-HCl 50 mM, NaCl 500 mM, pH 8.0]/glycerol 90:10, v/v). SMAc samples were then analyzed to evaluate the volume-distribution mean diameter 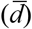 of SMAc particles as a function of the buffer pH and ionic strength (*I*).

**Table 2.** Potassium phosphate buffers (KP) used for the DLS analysis on SMAc solubility.

| Variable | Label | Buffer composition | pH | $I$ (M) |
| --- | --- | --- | --- | --- |
| pH | KP100 | potassium phosphate 100 mM | 6.0 | 0.112 |
|  |  |  | 7.0 | 0.176 |
|  |  |  | 8.0 | 0.272 |
|  |  |  | 9.0 | 0.297 |
| ionic strength | KP10 | potassium phosphate 10 mM | 8.0 | 0.027 |
|  | KP50 | potassium phosphate 50 mM | 8.0 | 0.136 |
|  | KPS50 | potassium phosphate 50 mM, NaCl 150 mM | 8.0 | 0.286 |
|  | KPS100 | potassium phosphate 100 mM, NaCl 150 mM | 8.0 | 0.422 |

### 2.5 Production of rhβ_2_AR-SMALPs

#### 2.5.1 Turbidimetric assays

Turbidimetric assays were performed at room temperature on a V-530 spectrophotometer (Jasco, Japan), using a detection wavelength of 400 nm and a QS quartz microcuvette with a 1 cm optical pathlength (Hellma Analytics, Germany). Wild-type (wt-) HEK293T membrane suspensions (2%, m/v) were prepared in SB500 and SB0 buffers (pH 8.0). SMAc working solutions (1%, m/v) were prepared by dilution with KP10 buffer (pH 8.0) starting from stock solutions (5%, m/m) in KP10 buffer (pH 8.0) for X25 and X30, in aqueous KOH (0.15 M) for C30. SMALP samples were prepared by mixing membrane suspensions and SMAc working solutions in a 1:1 (v/v) proportion, yielding a membrane concentration of 1% (m/v) and a membrane/polymer ratio of 2:1 (m/m). Turbidimetric measurements were carried out immediately after sample preparation and at 5 min intervals for 45 min. The residual turbidity of each membrane suspension after treatment with different SMAc co-polymers was then determined by comparing the optical density at 400 nm (OD_400_) with that of the same wt-HEK293T membrane suspension after 1:1 (v/v) dilution with KP10 buffer (pH 8.0).

#### 2.5.2 SEC analysis

The optimization of experimental conditions to maximize the yields of SMALP production was carried out by SEC analysis on a PU-2089plus HPLC system (Jasco, Japan) equipped with a UV-2077 UV-Vis detector (detection wavelength: 280 nm), an AS-2057 autosampler (injection volume: 200 μL) and a Superdex 200 10/300 GL column (GE Healthcare Life Sciences, UK; column volume: 23 mL). The influence of SMAc and salt concentrations on the yields of SMALP production was evaluated by varying the concentration of NaCl in the suspension and running buffers used during the preparation and purification of SMALPs. Running buffers RB500 (potassium phosphate 50 mM, NaCl 500 mM, pH 8.0) and RB0 (potassium phosphate 50 mM, pH 8.0) were therefore tested as mobile phases for the SEC runs (30 mL) at flow rates of 0.3–0.5 mL/min, depending on the initial backpressure. On the other hand, wt- or rhβ_2_AR-overexpressing HEK293T membrane suspensions (8%, m/v) were prepared in SB500, SB250 and SB0 buffers (Table 1). Finally, SMAc solutions were prepared in KP10 buffer (pH 8.0) at different concentrations ranging from 0.8 to 8% (m/v). SMALP samples were prepared by mixing membrane suspensions and SMAc solutions in a 1:1 (v/v) proportion, yielding a constant membrane concentration of 4% (m/v) and different membrane/polymer ratios ranging from 1:1 to 10:1 (m/m); samples were always annealed at 37 °C for 30 min under sonication and ultracentrifuged at 4 °C and 100000×*g* for 20 min before analysis.

#### 2.5.3 SEC–IMAC purification

Purification of rhβ_2_AR-SMALPs was achieved by two-dimensional liquid chromatography (2D-LC), in which SEC and immobilized metal affinity chromatography (IMAC) were combined through a switch valve (Figure 2). The same HPLC system used for SEC analysis was employed; a Superdex 200 10/300 GL column was used for SEC separation, while a Tricorn 5/20 column (GE Healthcare Life Sciences, UK; column volume: 0.5 mL) packed with a nickel(II)–nitrilotriacetic acid (Ni–NTA) agarose resin (Agarose Bead Technologies, USA) was used for IMAC. Running buffer RB0 and elution buffer EB (imidazole 200 mM in potassium phosphate 50 mM, pH 8.0) were used as mobile phases at a flow rate of 0.4 mL/min.

**Figure 2.**
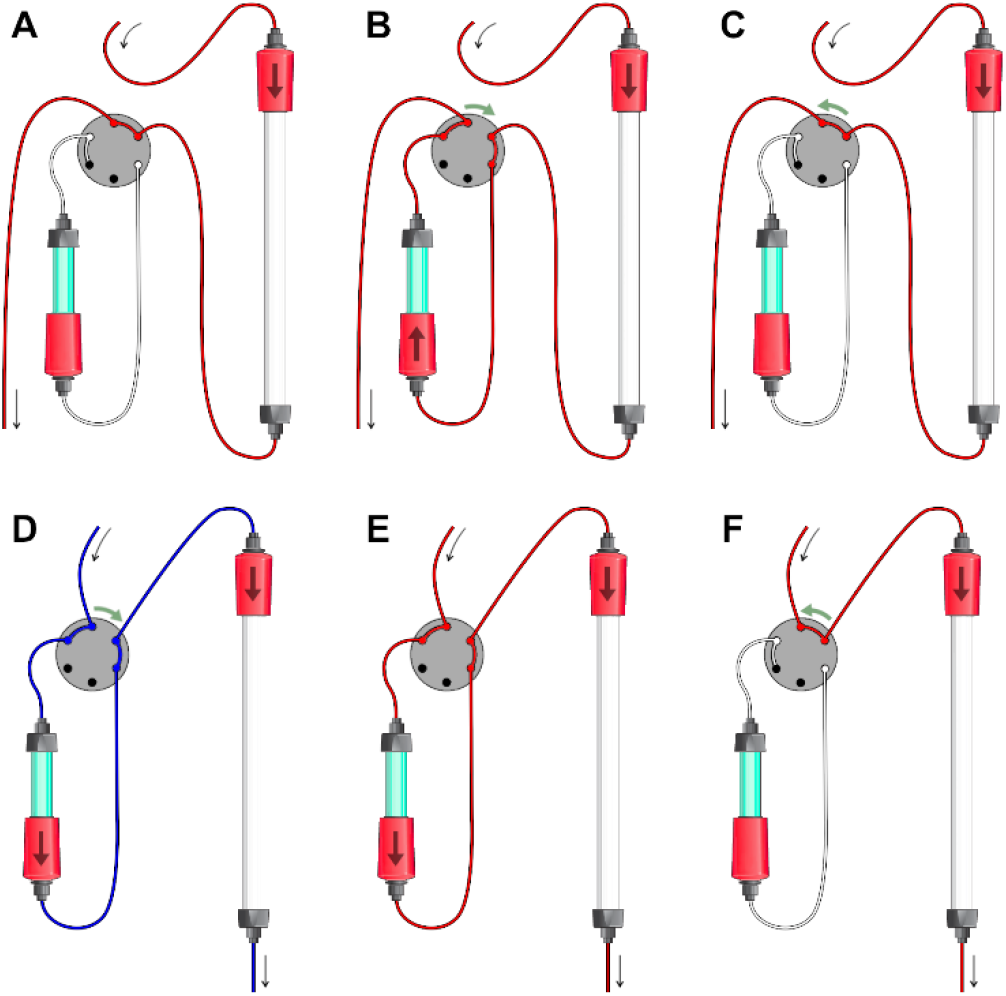
2D-LC setup for the SEC–IMAC purification of rhβ_2_AR-SMALPs in the loading step (**A:** 0–12 mL; **B:** 12–19 mL; **C:** 19– 30 mL) and in the elution step (**D:** 0–5 mL; **E:** 5–10 mL; **F:** 10–40 mL). The SEC (white) and IMAC (*cyan*) columns, as well as the RB0 (*red*) and EB (*blue*) buffers used as mobile phases, are depicted. Arrows indicate flow directions and valve operations at the beginning of each step.

Before use, the IMAC column was regenerated by flushing the resin sequentially with Na2EDTA (0.1 M in water) and with a fresh NiCl2 solution (0.1 M in water). The purification took place in two steps (Figure 2): (**a**) loading step (30 mL); (**b**) elution step (40 mL). During the loading step (Figure 2A–C), the IMAC column was connected in normal flow direction to the downstream end of the SEC column and the RB0 buffer was used as mobile phase in isocratic conditions. During the elution step (Figure 2D–F), the IMAC column was connected in reverse flow direction to the upstream end of the SEC column and a step gradient was used for the mobile phase: *i.e.*, EB buffer for the first 5 mL (Figure 2D), then RB0 buffer until the end of the run (Figure 2E–F). The IMAC column was excluded by default from the flow of the HPLC system, but the connection between the SEC and IMAC columns was devised with a 7000 two-position switching valve (Rheodyne, USA) in order to flush the IMAC column only at specific times during the runs: *i.e.*, in the volume range of SMALP elution during the loading stage (12–19 mL, Figure 2B) and at the beginning of the elution stage (0–10 mL, Figure 2D–E).

The SMALP sample was prepared by mixing a rhβ_2_AR-HEK293T membrane suspension (8% in SB0 buffer, m/v) and a SMAc solution (X30, 0.8% in KP10 buffer, m/v) in a 1:1 (v/v) proportion, yielding a membrane concentration of 4% (m/v) and a membrane/polymer ratio of 10:1 (m/m); the sample was annealed at 37 °C for 30 min under sonication and ultracentrifuged at 4 °C and 100000×*g* for 20 min before injection (volume: 200 μL). Purified rhβ_2_AR-SMALPs were collected during the elution stage, in the volume range (15–21.6 mL) over which the UV signal at 280 nm diverged significantly from the baseline signal recorded with a blank injection. The resulting 5.6-mL fraction was concentrated to 250 μL by ultrafiltration using Amicon Ultra-4 10K centrifugal filters (Merck, USA), then submitted to immunodetection.

#### 2.5.4 Immunodetection

The presence of rhβ_2_AR in the SEC–IMAC fraction was verified by dot-blot immunodetection. The concentrated SEC–IMAC fraction was blotted on two dry nitrocellulose membranes (Bio-Rad, USA) by applying 2 μL of each sample. Due to the low expected amount of rhβ_2_AR-SMALPs, in addition to concentration of samples, the samples obtained in SEC-IMAC were added by 2 μL aliquots on the same spot 25 times (total volume: 50 μL); after each application, the membranes were left to dry. As positive controls, protein solutions (2 μL) bearing appropriate peptide tags were used: *i.e.*, 6×His-tagged recombinant MSP1E3D1 (Sigma-Aldrich, Merck, USA) and FLAG-tagged recombinant mCardinal (kindly provided by Jakub Czapiński, Department of Biochemistry and Molecular Biology, Medical University of Lublin, Poland). The dot-blot membranes were blocked for 30 min with skim milk (3%, v/v) in TBS-T buffer (Tris-HCl 20 mM, NaCl 150 mM, pH 7.4 + Tween-20 0.1%, v/v), then incubated overnight at 4 °C with monoclonal primary antibodies directed against both peptide tags present on rhβ_2_AR: *i.e.*, anti-6×His and anti-FLAG M2 (Sigma-Aldrich, Merck, USA). The anti-mouse IgG, horseradish peroxidase (HRP)-linked rabbit antibody (Cell Signaling, USA) was used as a secondary antibody (30 min incubation at room temperature). The Westar Supernova substrate for enhanced chemiluminescence (ECL) detection (Cyanagen, Italy) was used to visualize the membranes; ECL was recorded using an Azure c400 western blot imaging system (Azure Biosystems, USA) with an exposure time of 1800 and 3600 s for membranes treated with anti-6×His and anti-FLAG M2 primary antibodies, respectively.

## 3 Results and discussion

### 3.1 Production of rhβ_2_AR

The purposely designed rhβ_2_AR form was devised in order to facilitate expression, isolation and detection once embedded into SMALPs. The cleavable ssHA signal sequence enhances the expression of functional receptors [23], while dual peptide tagging on both termini was chosen for redundancy reasons. The *N*-terminal FLAG tag allows for unambiguous immunodetection by exploiting the high selectivity and specificity of anti-FLAG M2 antibodies; on the other hand, the *C*-terminal double His tag (10×His, 12-residue spacer. 6×His) was inserted to ensure a more stable interaction with the Ni–NTA resin used for the downstream purification of SMALPs by IMAC [24]. C-terminal His tags were preceded by the TEV site for subsequent cleavage in planned future experiments.

A prerequisite for the selection of the expression system was the ability to provide a native membrane environment around rhβ_2_AR during SMALP formation in the form of an annular lipid shell. The human embryonic kidney 293T cell line (HEK293T) was chosen for its robustness in culture and ease of transfection, high efficiency of protein production and generally for being a proven recombinant receptor platform [25]. Transient expression using cationic transfection compound was chosen on the basis of rapidity of the process and high protein expression levels [26,27].

The successful expression of rhβ_2_AR was confirmed by dot-blot immunodetection. Since the recombinant protein is flanked by two peptide tags on both the *N*- and *C*-termini, detection of both of these tags provides information on the complete translation of the protein (Figure 3). Out of the obtained clones, clone 3 was used for further processing due to the most uniform expression of both tags. The correct nucleotide sequence encoding for rhβ_2_AR in the plasmid was confirmed by Sanger sequencing.

**Figure 3.**
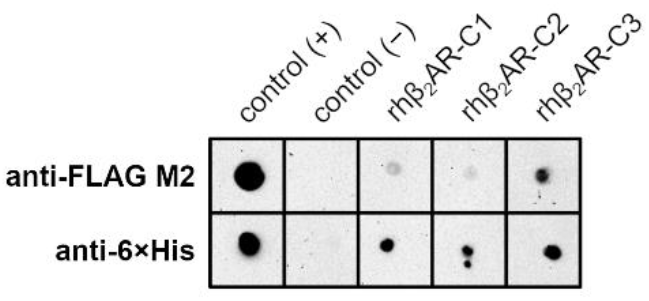
Dot-blot immunodetection of rhβ_2_AR expression. Positive control: mCardinal-FLAG and MSP1E3D1-6×His, respectively. Negative control: raw membrane suspension from wt-HEK293T cells. rhβ_2_AR-C1, -C2, -C3: raw membrane suspensions from discrete clones of rhβ_2_AR-HEK293T cells.

### 3.2 Solubility of SMAc co-polymers

The amphiphilic character of SMAc co-polymers is one of the most important features of the SMALP technology. The ideal SMAc co-polymer for a specific application should provide the best balance between the lipophilicity of Sty moieties, which mediate the affinity toward the target cell membranes, and the hydrophilicity of MAc moieties, which regulate the stabilization of SMALPs in water [28]. Therefore, the first important step in the production of SMALPs is the selection of the SMAc co-polymer with the most suitable properties in terms of MP solubilization from the cell line chosen for the overexpression of the GPCR under investigation, including solubility in water.

The solubility of SMAc co-polymers in aqueous media is strongly influenced by several structural and environmental factors, such as Sty/MAc ratio, molecular weight, presence of counter-ions, as well as pH and ionic strength of the solvent [19,29–31]. The degree of protonation of carboxyl groups of SMAc increases as the pH of the solution decreases, enhancing the hydrophobicity of the co-polymer and promoting aggregation (Figure 4A). As a general trend, the particle size of the commercial co-polymer X30 (MAc content: 30%, n/n; 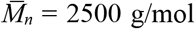; 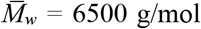) was found to be smaller than that of the commercial co-polymer X25 (MAc content: 25%, n/n; 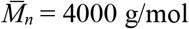; 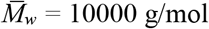) and, to a much greater extent, that of the in-house synthesized co-polymer C30 (MAc content: 30%, n/n; 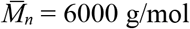; 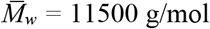; details on the synthesis of C30 are reported in the Supplementary Information).

**Figure 4.**
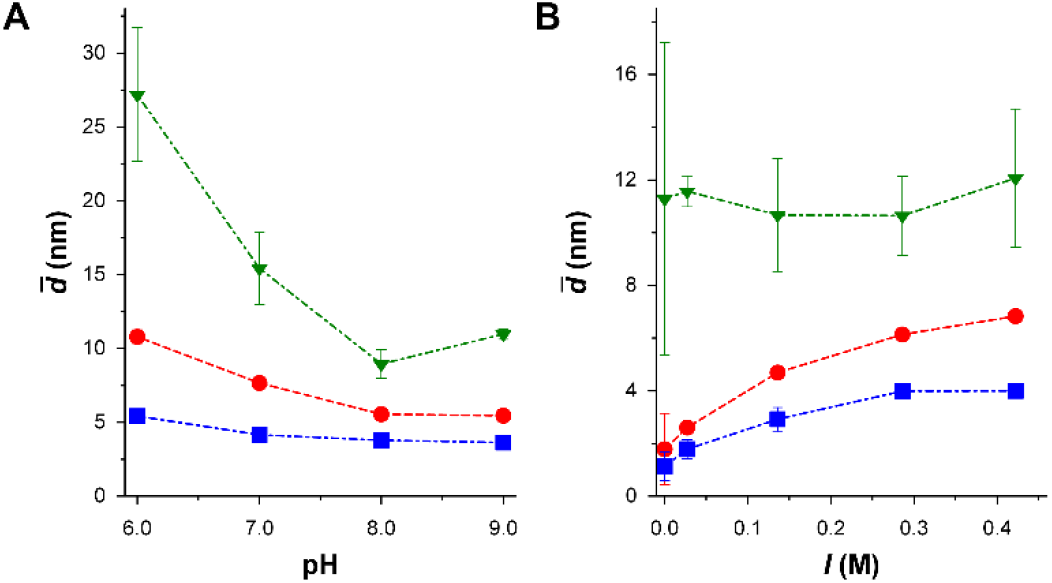
DLS measurements on SMAc co-polymers X25 (*circles*), X30 (*squares*) and C30 (*triangles*). SMAc particle sizes (*n* = 3) are reported as a function of pH at constant buffer composition (**A**) and ionic strenght at constant buffer pH (**B**). Buffer compositions are given in Table 2.

Presence of counter-ions in the starting SMAc batch was also a critical factor: X25 and X30 were supplied as potassium salts in highly concentrated aqueous solutions (>20%, m/m), whereas C30 was produced as a free acid in solid form and subsequently dissolved in the presence of KOH (0.15 M). Consequently, while the particle size of C30 was constantly larger irrespective of the ionic strength of the solvent, a significant salting-out effect could be observed for X25 and X30, showing that the potassium salts of SMAc co-polymers tend to aggregate in saline solutions (Figure 4B).

Based on these observations, pure water and diluted buffers (such as KP10) are more appropriate solvents in the preparation of SMAc stock solutions in order to achieve maximum solubility for X25 and X30. However, buffered stock solutions also provide pH control, ensuring an optimal ionization of the co-polymer after mixing with cell membranes, which are typically suspended in SB500 buffer (pH 8.0) according to the standard protocol for SMALP preparation. Therefore, SMAc stock solutions for the subsequent biophysical assays on SMALP formation were prepared in KP10 buffer (pH 8.0).

### 3.3 Production of rhβ_2_AR-SMALPs

#### 3.3.1 Turbidimetric assays

The treatment of cell membranes with SMAc causes a decrease in the turbidity of membrane suspensions due to the formation of SMALPs; therefore, turbidimetric assays are a very practical analytical tool for fast qualitative evaluations on the kinetics and yield of this process, even though their accuracy for quantitative purposes is questionable. The turbidimetric profiles of wt-HEK293T membrane suspensions treated with SMAc (Figure 5) showed that the process of SMALP formation is influenced by the Sty/MAc ratio of the co-polymer and by the ionic strength of the solvent. These factors were shown to influence the size of polymer particles in aqueous media as well: this observation suggests a strong link between the water solubility of SMAc and its solubilization properties toward membranes and membrane proteins.

**Figure 5.**
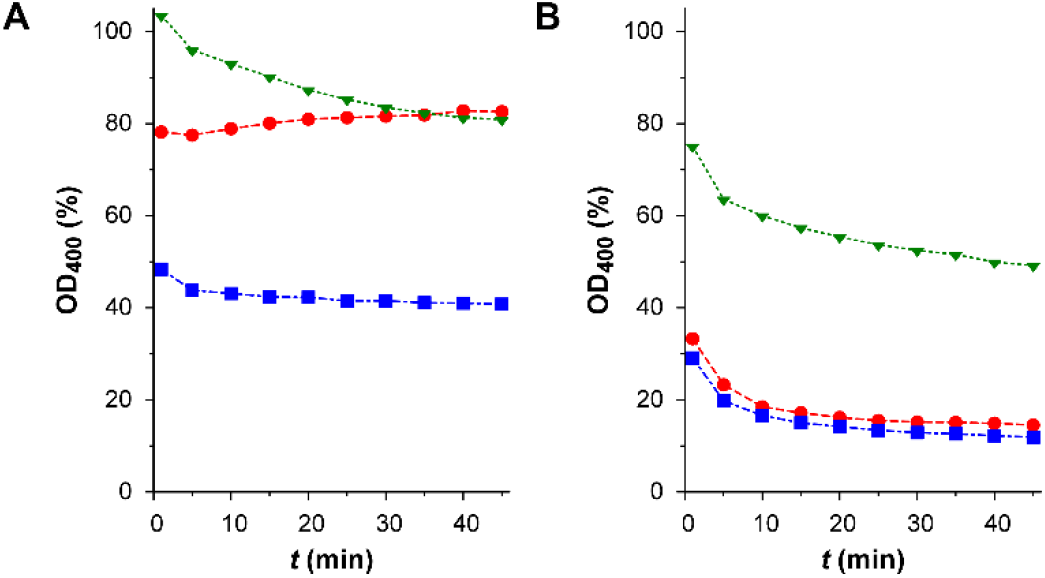
Turbidimetric profiles of SMALP formation for wt-HEK293T membrane suspensions (2%, m/v) mixed 1:1 (v/v) with different SMAc co-polymers (1% in KP10 buffer, pH 8.0, m/v): X25 (*circles*), X30 (*squares*) and C30 (*triangles*). (**A**) Membranes suspended in SB500 buffer. (**B**) Membranes suspended in SB0 buffer.

Overall, the synthesized co-polymer C30 always elicited a lower reduction in turbidity compared to the commercial co-polymers X25 and X30 due to its lower solubility, which was earlier highlighted by DLS measurements (Figure 4). The high NaCl concentration (500 mM) of the SB500 buffer used to suspend cell membranes apparently interfered with the SMALP formation properties of X25, whose MAc content is lower compared to C30 and X30, resulting in a slight increase of turbidity with time (Figure 5A). When the salt concentration in the suspension buffer was reduced (SB0), the membrane solubilization properties of X25 improved dramatically, nearly matching the turbidimetric profile obtained with X30 (Figure 5B). Considering the results of DLS analysis and turbidimetric assays, X30 emerged as the SMAc co-polymer with the highest potential to provide better yields in the preparation of SMALPs with HEK293T membranes. Moreover, the strong influence of ionic strength on SMAc prompted additional investigations on the use of saline solutions in other steps of the SMALP protocol and their effect on the efficiency of SMALP preparation.

#### 3.3.2 SEC analysis on SMAc co-polymers

After SMALPs are produced by treatment of cell membranes with SMAc, SEC fractionation and affinity-based purification are typically required to purify SMALPs embedding the desired GPCR and isolate them from the whole pool of SMALPs embedding other MPs [15,32]. According to standard SEC protocols, a major source of salts is the running buffer used for the fractionation of SMALPs: the standard SEC buffer (RB500) contains 500 mM of NaCl. The SEC analysis on X30 using running buffers of different NaCl content (Supplementary Figure S4A) showed that SMAc tends to form larger aggregates at higher ionic strength (RB500), resulting in broader peaks and an elution range extending beyond the exclusion limit of the SEC column, which is a warning sign of the occurrence of hydrophobic interactions between the co-polymer and the column matrix. At a lower ionic strength (RB0), X30 showed a better profile, as the co-polymer eluted as a narrower peak within the exclusion limit. This behavior also affects the SEC profiles of SMALP samples: the higher intensity of overlapped peaks in the elution range of SMALPs is due to the concurrent co-elution of SMAc aggregates (Figure 6A). Because of peak broadening, higher ionic strengths of the running buffer have a negative effect on the purity of SMALP fractions during SEC fractionation.

**Figure 6.**
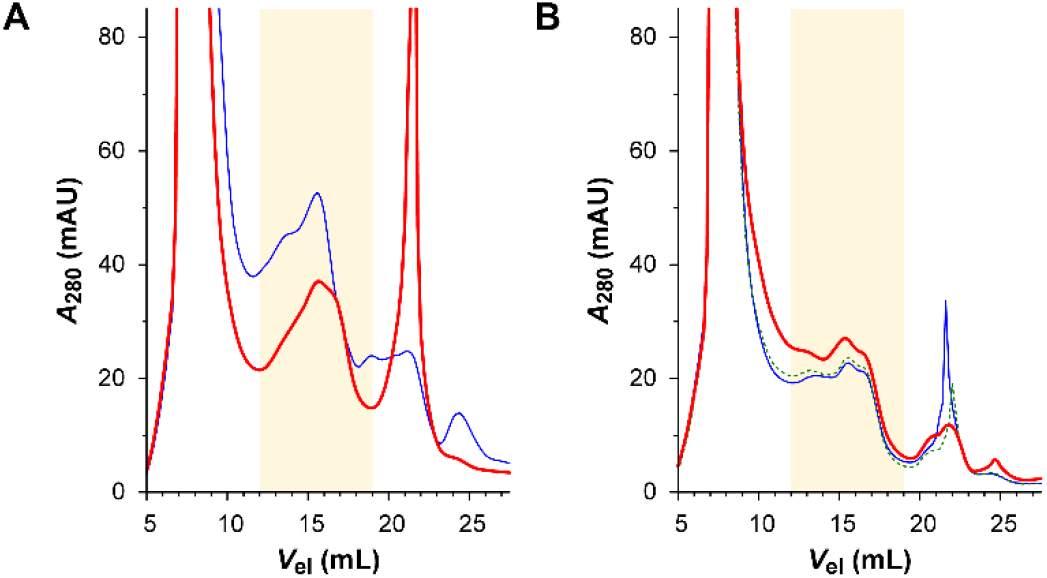
Chromatograms (280 nm) of SMALP samples obtained from 1:1 (v/v) mixtures of wt-HEK293T membrane suspensions (8%, m/v) and SMAc co-polymer X30 in KP10 buffer (pH 8.0), as obtained by SEC analysis on a Superdex 200 10/300 GL column; the range of SMALP elution (12–19 mL) is highlighted. (**A**) Effect of salt concentration in SEC buffers on SMALP purification. Samples were prepared using X30 4% (m/v) and membranes suspended in SB500 buffer; SEC runs were performed using RB500 (*thin*) and RB0 (*bold*) buffers as mobile phases. (**B**) Effect of salt concentration in SMALP buffer on SMALP formation. Samples were prepared using X30 0.8% (m/v) and membranes suspended in SB500 (*thin*), SB250 (*dotted*) or SB0 (*bold*) buffers; SEC runs were performed using RB0 buffer as mobile phase.

Another major source of salts is the buffer used to suspend membranes; in agreement with results from turbidimetric assays, the intensity of SEC peaks relative to SMALP fractions was higher when the NaCl concentration of the suspension buffer was reduced, indicating a larger yield of SMALP formation. This behavior was observed with both wt-(Figure 6B) and rhβ_2_AR-HEK293T (Figure 7A) membranes treated with X30, suggesting that the protein density on the membrane surface does not influence the SMALP formation efficiency of SMAc. Therefore, better yields in the preparation of SMALPs with rhβ_2_AR-HEK293T membranes was achieved by lowering the ionic strenght, e.g. by removing NaCl from all buffers.

**Figure 7.**
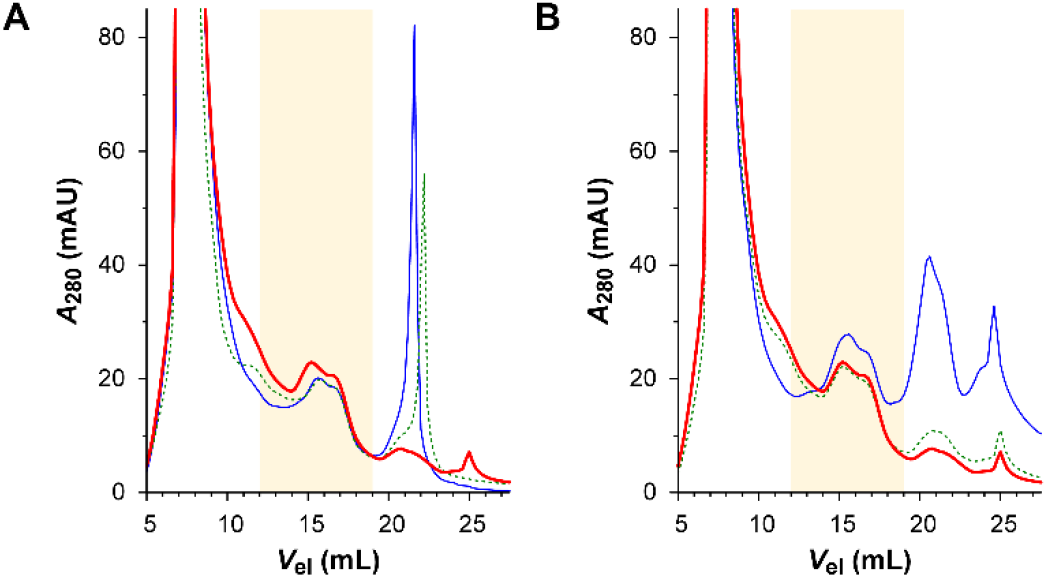
Chromatograms (280 nm) of SMALP samples obtained from 1:1 (v/v) mixtures of rhβ_2_AR-HEK293T membrane suspensions (8%, m/v) and SMAc co-polymer X30 in KP10 buffer (pH 8.0), as obtained by SEC analysis on a Superdex 200 10/300 GL column using RB0 buffer as mobile phase; the range of SMALP elution (12–19 mL) is highlighted. (**A**) Effect of salt concentration in SMALP buffer on SMALP formation. Samples were prepared using X30 0.8% (m/v) and membranes suspended in SB500 (*thin*), SB250 (*dotted*) or SB0 (*bold*) buffers. (**B**) Effect of SMAc concentration on SMALP formation. Samples were prepared using X30 8% (m/v; *thin*), 2% (m/v; *dotted*) or 0.8% (m/v; *bold*) and membranes suspended in SB0 buffer.

The effect of ionic strength on SMALP formation has been already evaluated and thoroughly discussed in the framework of an optimization study for the isolation of MPs from *Escherichia coli* cells [17], where the solubilization of the KcsA potassium channel was found to be gradually enhanced by the presence of NaCl concentrations up to 300 mM, then inhibited by higher concentrations. This behavior was observed using SMAc in acid form as obtained by hydrolysis of the commercial SMAn co-polymer Xiran SZ 30010, which is supplied by Polyscope as a powder; on the other hand, our results were obtained with Polyscope’s X30 (Xiran SL 30010 P20), which is a solution of SMAc potassium salt in water. The different effect of ionic strength is therefore explained by the different protonation state of the SMAc co-polymer in the two studies, further highlighting the need for a rigorous control of all the experimental factors influencing the efficiency of the SMALP formation process.

An additional factor to evaluate in the optimization of the SMALP protocol is the concentration of SMAc. Increasing the concentration of X30 in the stock solution from 0.8% to 2% (m/v) did not lead to a parallel increase in SMALP formation, i.e. the intensity of SEC peaks in the range of SMALP elution did not increase (Figure 7B). Therefore, a membrane/polymer ratio of 10:1 (w/w) is sufficient to achieve the maximum yield of SMALP formation while limiting the interference of free SMAc. Indeed, a partial co-elution of free co-polymer and SMALPs occurred when higher concentrations of SMAc were used, as observed when X30 stock solutions were prepared at 8% (m/v) (Figure S4C in the Supplementary Information).

#### 3.3.3 SEC–IMAC purification

The purification of SMALPs is achieved by a two-step procedure: the separation of SMALPs from larger membrane fractions and free SMAc is achieved by SEC, while the isolation of SMALPs embedding the membrane protein of interest from the rest of SMALPs is achieved by affinity chromatography, e.g. IMAC if the membrane protein bears a poly-His tag, as in the case of the rhβ_2_AR customly designed for this study. SEC fractionation is preferably performed before IMAC isolation to avoid interference from membrane fractions and free SMAc in the affinity interaction between immobilized metals and His-tagged SMALPs [33,34]; preliminary tests performed during our study showed that SMAc can lead to a ~25% loss in the retention of soluble His-tagged proteins onto the Ni–NTA resin (data not shown). A 2D-LC combination of SEC fractionation and IMAC isolation was therefore designed (Figure 2). In this experimental setup, the rhβ_2_AR-SMALP fraction isolated by IMAC is eluted with imidazole and directly processed through a second SEC fractionation, in order to purify SMALPs from imidazole and enable real-time monitoring by UV detection throughout the process.

Based on the outcomes of optimization studies on SMALP production, rhβ_2_AR-SMALPs were prepared with 1:1 (v/v) mixtures of rhβ_2_AR-HEK293T membranes suspensions (8% in SB0 buffer, pH 8.0, m/v) and X30 (0.8% in KP10 buffer, pH 8.0, m/v). When the IMAC column is connected, the eluate from the SEC column is transferred to the IMAC column, which retains His-tagged rhβ_2_AR-SMALPs while other SMALPs flow through without interacting with the Ni–NTA resin; therefore, the resulting chromatographic profile (bold line in Figure 8A) accounts for the IMAC-unretained material. The two flow discontinuities at 12 and 19 mL in the SEC chromatogram (Figure 8A) are the effect of valve switching to connect the IMAC column located downstream to the SEC column when SMALPs elute (12–19 mL).

**Figure 8.**
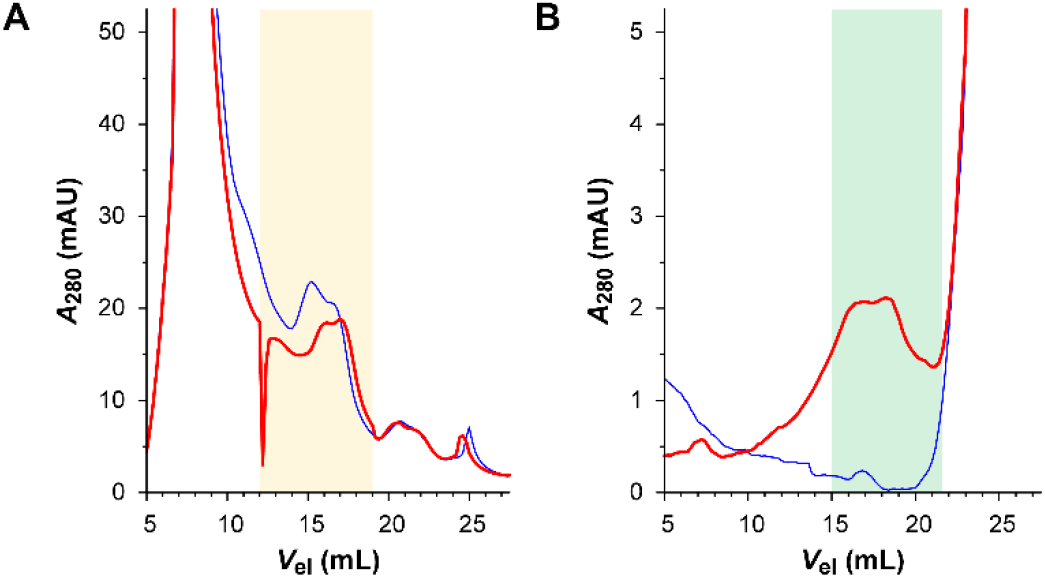
SEC–IMAC purification of rhβ_2_AR-SMALPs, as obtained with a Superdex 200 10/300 GL column coupled with a Tricorn 5/20 packed with Ni–NTA resin. (**A**) SEC chromatograms (280 nm) of SMALP samples obtained from a 1:1 (v/v) mixture of a rhβ_2_AR-HEK293T membrane suspension (8% in SB0 buffer, m/v) and SMAc co-polymer X30 (0.8% in KP10 buffer, pH 8.0, m/v), without treatment with IMAC (*thin*) and in the loading step of the SEC–IMAC purification (*bold*); the range of SMALP elution (12– 19 mL) is highlighted. (**B**) SEC chromatogram (280 nm) of purified rhβ_2_AR-SMALPs in the elution stage of the SEC–IMAC purification (*bold*) compared to the baseline signal recorded with a blank injection (*thin*); the range of SMALP collection (15–21.6 mL) is highlighted.

Comparing this profile with the chromatogram of a SEC separation of the same sample (thin line in Figure 8A), a clear difference in shape and intensity of the SMALP peak is observed. This difference can be interpreted as the result of the retention of the His-tagged rhβ_2_AR-SMALPs onto the the IMAC column, which is confirmed by the SEC analysis of the material eluted from the IMAC column after treatment with imidazole (Figure 8B).

Interestingly, the SEC profile of isolated rhβ_2_AR-SMALPs (Figure 8B) retained the typical twin-peaked shape of SMALPs; moreover, the designed 2D-LC setup successfully allowed the purification of rhβ_2_AR-SMALPs from imidazole. The intensity of the SEC peak clearly indicates that the net amount of purified rhβ_2_AR-SMALPs is small; before immunodetection, the fraction was therefore concentrated (~22×) by ultrafiltration. Dot-blot immunodetection (Figure 9) showed the presence of both tags incorporated into rhβ_2_AR in both raw membrane preparations and mix of thereof with SMAc (Figure 9). Albeit at a low concentration, the SEC-IMAC elution fraction contained *de facto* purified rhβ_2_AR-SMALPs. Due to the low amount, multiple depositions of the sample and long exposition times were required to qualitatively detect rhβ_2_AR, but the assessment of further elution fractions confirmed the correct identification of the SEC peak.

**Figure 9.**
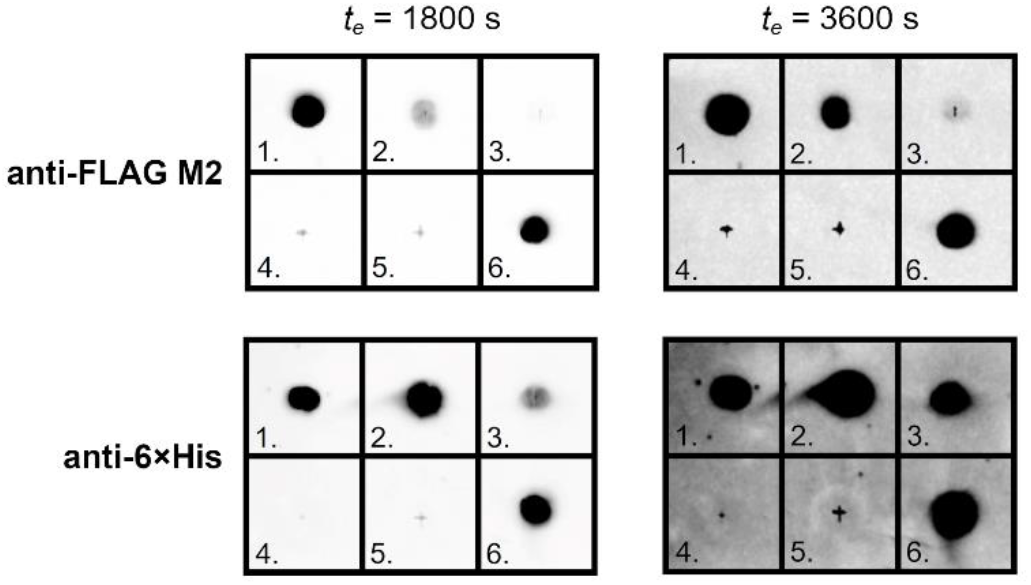
Results of dot-blot immunodetection. Samples used for the blot: **1.** rhβ_2_AR-HEK293T raw membrane suspension (120 mg/mL); **2.** 1:1 (v/v) mixture of a rhβ_2_AR-HEK293T membrane suspension (8% in SB0 buffer, m/v) and SMAc co-polymer X30 (0.8% in KP10 buffer, pH 8.0, m/v); **3.** Purified rhβ_2_AR-SMALPs (15–21.6 mL range); **4.** SEC-IMAC elution fraction (21.6–23.6 mL range); **5.** SEC-IMAC elution fraction (23.6–25.6 mL range); **6.** Positive control: mCardinal-FLAG and MSP1E3D1-6×His, respectively. Samples 3–5 were spotted 25×2 μL on the membrane. Exposition time for collecting ECL specified above.

## 4 Conclusions

The present article reports on our efforts aiming at isolating a recombinant form of the human β_2_-adrenergic receptor from purposely engineered HEK293T cells using the SMALP technology. The inherent limitations of using a low-yielding mammalian expression system prompted us to focus our attention on optimizing the yields of SMALP production by tuning several experimental parameters. In addition to proper recombinant vector design, composition of the employed SMAc co-polymer, the pH and ionic strength of solubilization buffers and the composition of running buffers appeared to play important role for purification of rhβ_2_AR-SMALPs. In particular, the ionic strength of buffers was found to be a critical factor governing the solubility of SMAc in water and the behavior of SMAc and SMALPs in the SEC fractionation and IMAC isolation steps of SMALP production. Moreover, a 2D-LC combination of SEC and IMAC was designed to monitor the purification process in real time while losses of purified rhβ_2_AR-SMALPs due to fraction collection were minimal. The purification of SMALP-embedded rhβ_2_AR from an overexpressing HEK293T cell line proved to be a challenge and highlighted the need for a careful optimization of some key experimental factors during the production of SMALPs, such as the ionic strength of buffers. This need might be easily overlooked when large amounts of biological material are available, but it should always be considered whenever the SMALP technology is applied to isolate GPCRs from low-yielding expression systems, such as human cell lines, in order to maintain mammalian PTMs or preserve the native lipid environment of GPCRs for functional studies.

## Abbreviations

2D-LC: two-dimensional liquid chromatography
AIBN: 2,2’-azobisisobutyronitrile
ATR-FTIR: attenuated total reflectance Fourier-transform infrared
DLS: dynamic light scattering
DMEM: Dulbecco’s Modified Eagle Medium
EB: elution buffer
ECL: enhanced chemiluminescence
FBS: fetal bovine serum
GPCR: G protein-coupled receptor
HPLC: high-performance liquid chromatography
HRP: horseradish peroxidase
IMAC: immobilized metal affinity chromatography
KP: potassium phosphate buffer
MAc: maleic acid
MAn: maleic anhydride
MP: membrane protein
Ni–NTA: nickel(II)–nitrilotriacetic acid
NMR: nuclear magnetic resonance
PCR: polymerase chain reaction
PTM: post-translational modification
RB: running buffer
rhβ_2_AR: recombinant human β_2_-adrenergic receptor
SB: suspension buffer
SEC: size-exclusion chromatography
SMAc: styrene–maleic acid co-polymer
SMAn: styrene–maleic anhydride co-polymer
SMALP: styrene–maleic acid lipid particle
ssHA: hemagglutinin signal sequence
Sty: styrene
TEV: tobacco etch virus
THF: tetrahydrofuran

## Acknowledgements

Dr. Stefan Scheidelaar (Polyscope Polymers BV, Geleen, Netherlands) for kindly providing commercial SMAc co-polymers; Dr. Jakub Czapiński (Department of Biochemistry and Molecular Biology, Medical University of Lublin, Poland) for fruitful discussion on the production of the plasmid construct and for kindly providing the positive control for the immunodetection of FLAG-tagged proteins; Ms. Alessia Berardinelli and Ms. Sara Zinato (Department of Pharmacy and Biotechnology, University of Bologna, Italy) for technical assistance during the production of the plasmid construct; Dr. Piotr Drączkowski (Chair and Department of Synthesis and Chemical Technology of Pharmaceutical Substances, Medical University of Lublin, Poland) for fruitful discussion on MP solubilization techniques.

## Author contributions

**Daniele Tedesco:** Conceptualization, Methodology, Investigation, Validation, Data curation, Visualization, Writing - original draft preparation. **Maciej Maj:** Conceptualization, Methodology, Investigation, Validation, Data curation, Visualization, Writing - original draft preparation. **Paulina Malarczyk:**Investigation, Validation. **Andrea Cingolani:** Investigation, Validation. **Mirko Zaffagnini:** Methodology, Resources. **Artur Wnorowski:**Methodology. **Tiziana Benelli:** Methodology, Resources, Supervision. **Rita Mazzoni:**Methodology, Resources, Supervision. **Manuela Bartolini:**Conceptualization, Funding acquisition, Resources, Project administration, Supervision, Writing - review & editing. **Krzysztof Jóźwiak:** Conceptualization, Funding acquisition, Resources, Project administration, Supervision, Writing - review & editing.

## Supplementary material

Details on the synthesis and characterization of SMAc co-polymer C30 (including NMR and ATR-FTIR spectra); SEC chromatograms of SMAc co-polymer X30.

