## Supplementary Material for "Application of the SMALP technology to the isolation of GPCRs from low-yielding cell lines"

### S1 Synthesis and characterization of SMAc co-polymer C30

The synthesis of N30 by co-polymerization of Sty with MAn (Sty/MAn ratio 8:1 n/n) was carried out in THF solution under radical conditions in the presence of AIBN (4%, n/n) as thermal initiator, achieving a yield of 35%. The occurrence of polymerization involving the double bonds was confirmed by $^{1}$H-NMR analysis, which showed the lack of the resonances of vinylic protons of MAn and Sty (around 7.1, 5.7 and 5.2 ppm respectively). Furthermore, a new signal appeared in the 1–2 ppm region belonging to the aliphatic main chain. The final molar composition of N30 was assessed by $^{13}$C-NMR by comparing the integrated peak areas of MAn carbonyl carbon atoms, located at around 173 ppm, to those related to the aromatic carbon atoms of Sty co-units in the range 150–120 ppm (Figure S1). The pre-hydrolyzed copolymer showed a MAn content of 30% (Sty/MAn ratio: 2.3:1, n/n).

N30 was then hydrolyzed to C30 in alkaline solution [1]; the reaction was boosted by subsequent additions of NaOH and by increasing the reaction time (5 days). The ATR-FTIR analysis of the obtained white solid (Figure S2) confirmed the presence of the SMAc co-polymer: the absence of the signals at 1855 and 1775 cm$^{-1}$ related to the C=O stretching of MAn (Figure S3), and the presence of the signal at 1565 cm$^{-1}$ related to the COO$^{-}$ stretching of maleic acid (MAc) reasonably led us to assume that the hydrolytic cleavage was taken to completion. On the other hand, the $^{13}$C-NMR spectrum showed a downfield shift of the resonances of the carbonyl carbon atoms. The integration of the $^{13}$C-NMR spectrum confirmed that the Sty/MAc ratio was maintained after hydrolysis. The average molecular weights of C30 were assessed by SEC analysis in THF solution at 25 °C ($\bar{M}_n$ = 6000 g/mol; $\bar{M}_w$ = 11500 g/mol).

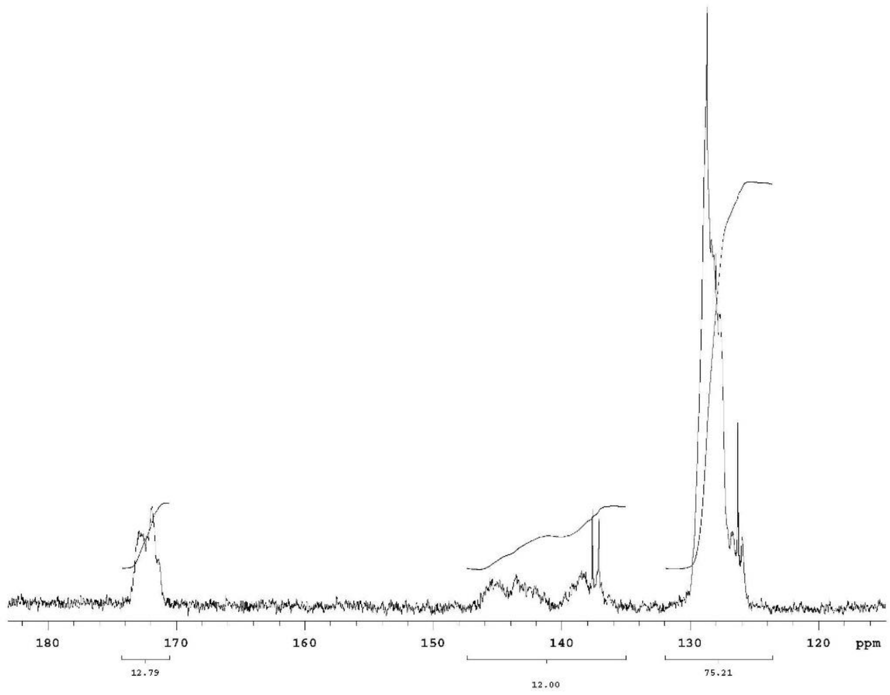


**Figure S1.** $^{13}$C-NMR spectrum of SMAn co-polymer N30 [solvent: $Cr(acac)_3$ 0.1 M in acetone-$d_6$].

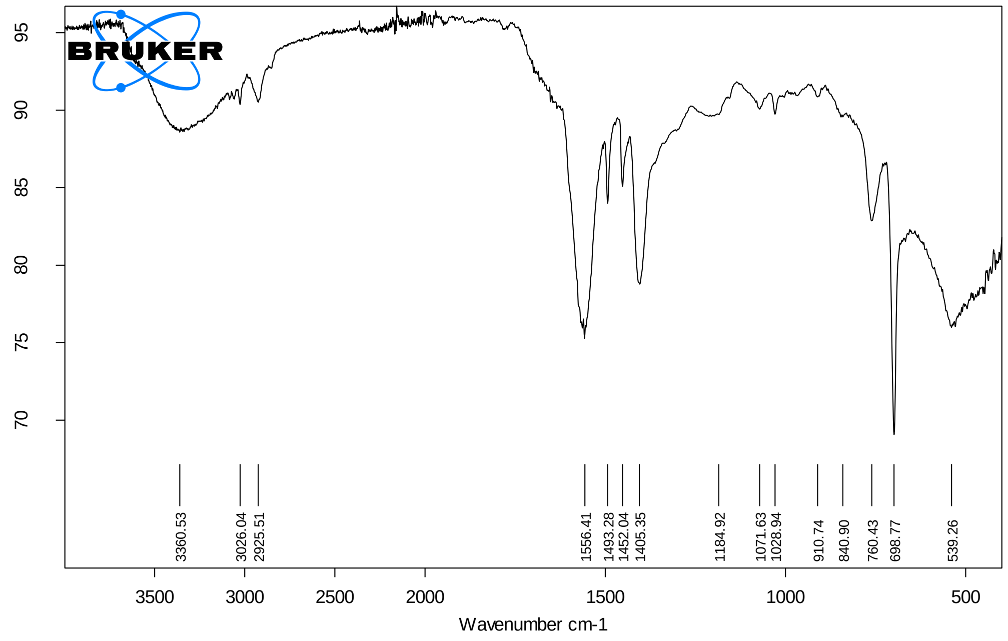


**Figure S2.** ATR-FTIR spectrum of SMAc co-polymer C30.

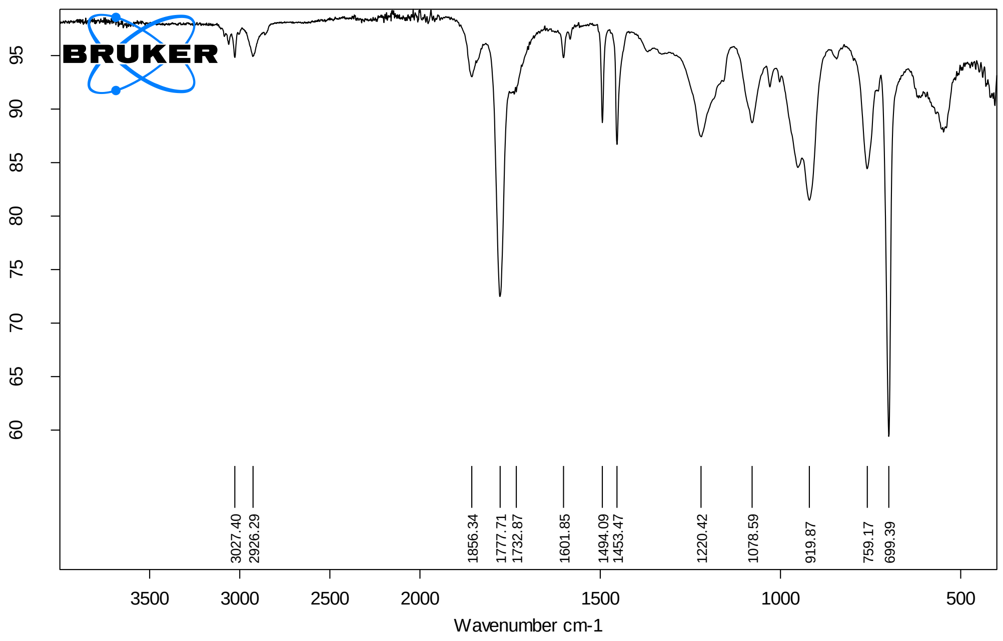


**Figure S3.** ATR-FTIR spectrum of SMAn co-polymer N30.

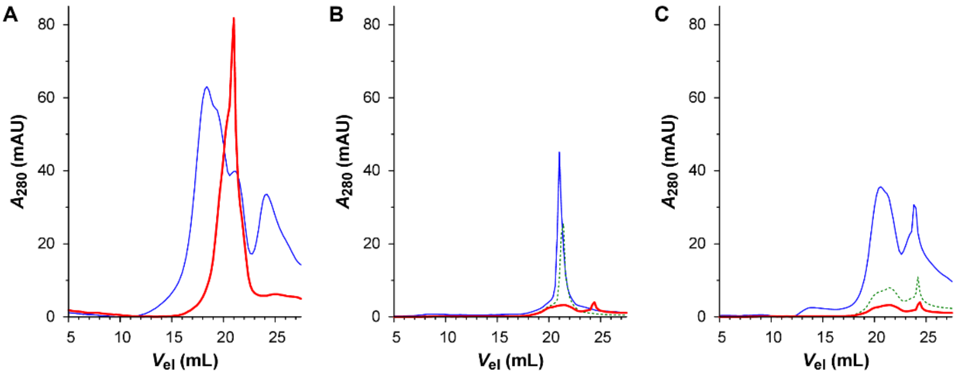


**Figure S4.** Chromatograms (280 nm) of SMAc co-polymer X30 in KP10 buffer (pH 8.0) mixed 1:1 (v/v) with SMALP buffers, as obtained by SEC analysis on a Superdex 200 10/300 GL column. (**A**) Samples were prepared with X30 4% (m/v) and SB500 buffer; SEC runs were performed using RB500 (*thin*) and RB0 (*bold*) buffers as mobile phases. (**B**) Samples were prepared with X30 0.8% (m/v) and SB500 (*thin*), SB250 (*dotted*) or SB0 (*bold*) buffers; SEC runs were performed using RB0 buffer as mobile phase. (**C**) Samples were prepared with X30 8% (m/v; *thin*), 2% (m/v; *dotted*) or 0.8% (m/v; *bold*) and membranes suspended in SB0 buffer; SEC runs were performed using RB0 buffer as mobile phase.
